# Landscape-scale predictors of mammal abundance and species richness across an extensive Queensland tropical savannas gradient

**DOI:** 10.1101/2025.09.22.677950

**Authors:** A.S. Kutt, H.S. Fraser

**Author notes:** **Declaration of funding:** No funding was received to prepare this manuscript. **Data availability:** The data will be published on *figshare* on acceptance of the manuscript for publication.

## Abstract

The small mammals in the tropical savannas of northern Australia, have undergone a degree of change in recent decades, best documented in the Northern Territory. Data is limited from northern Queensland and though the same trends are assumed, the topographic and climatic features differ substantially. In this study we examined data systematically collected from 725 sites between 1998-2012 in three bioregions representing a climatic gradient: from semi-arid to monsoon tropical savannas. We investigated via information-theoretic models and model averaging, the relationship between five mammal groupings and three landscape variables (fractional cover green, elevation and vegetation diversity) to elucidate any consistent or different patterns in the mammal fauna. Key patterns included relationships with increasing elevation (critical weight range species richness positively associated with elevation, rodent species richness negatively associated), increasing rodent and dasyurid species richness with vegetation diversity, and lower macropod and dasyurids abundance with increasing fractional cover green. These relationships underscore a need to consider mammal conservation in Queensland with more nuance than in the more topographically inert Northern Territory. Management strategies need to be more attuned to taxonomic and regional differences, to prevent perverse outcomes.

## INTRODUCTION

The tropical and sub-tropical savannas of northern Australia have a mammal fauna that is under distress, due to a range of long-acting threatening processes (Wallach and Lundgren 2025). The focus of most mammal research is on unpicking different interacting correlates of causation, such as fire and predation (Doherty *et al*. 2023). Furthermore, a major conservation approach has been to place mammals in safe havens free of threats (Legge *et al*. 2018) though there is an increasing focus on finding locations in the landscape that mammals can persist, where the threats are less or absent (Brewster *et al*. 2024).

Northern Australian tropical savannas are much more diverse than is often characterised (i.e., vast, limited variation, unbounded), given most of the descriptions of these landscapes are viewed through a Northern Territory lens (Woinarski *et al*. 2005). Though there is general correspondence of the nature of tropical savannas across the north, there is much more complex topographic, geological and condition variation in the eastern and western Australian expressions of tropical savannas. For example, the extensive climatic banding that features south to north in the Top End (Woinarski *et al*. 1999), is less apparent in Queensland due to the presence of the Great Dividing Range (which is significant topographical feature), and climate driven by both the monsoon and easterly south-easterly trade winds.

In this short study, we reviewed a large dataset of systematic fauna surveys in north-eastern Queensland to investigate the relationships broad landscape-scale variables and mammal species richness and abundance, across three bioregions, representing a gradient from the more southern, semi-arid tropical savannas to the more monsoon dominated locations. Understanding the more specific differences in tropical savanna biota in northern Australia, is important to more effectively consider the management of threatened species, threatening process, and conservation planning, especially as climate change more strongly influences the weather patterns in these biomes.

## METHODS

The mammal surveys were conducted across three Queensland tropical savanna bioregions that encompass part of the Great Dividing Range in Eastern Australia; the Desert Uplands, the Einasleigh Uplands, and Cape York Peninsula (Fig. 1). The tropical savannas of northern Australia generally occur in across a region demarked by line running from Townsville on the east coast to Broome on the west coast. Though collectively, they fall within the tropical savannas (which is broadly defined by distinctive rainfall seasonality with most of the rainfall occurring in between December and April), the three bioregions represent a gradient from the south to north, and from sub-tropical to monsoon tropical environments. All survey sites were vegetation dominated by mixed *Eucalyptus* (ironbark), *Corymbia* (bloodwood) and *Melaleuca* (ti tree) species, grasslands and occasional closed forests in riparian area.

**Figure 1.**
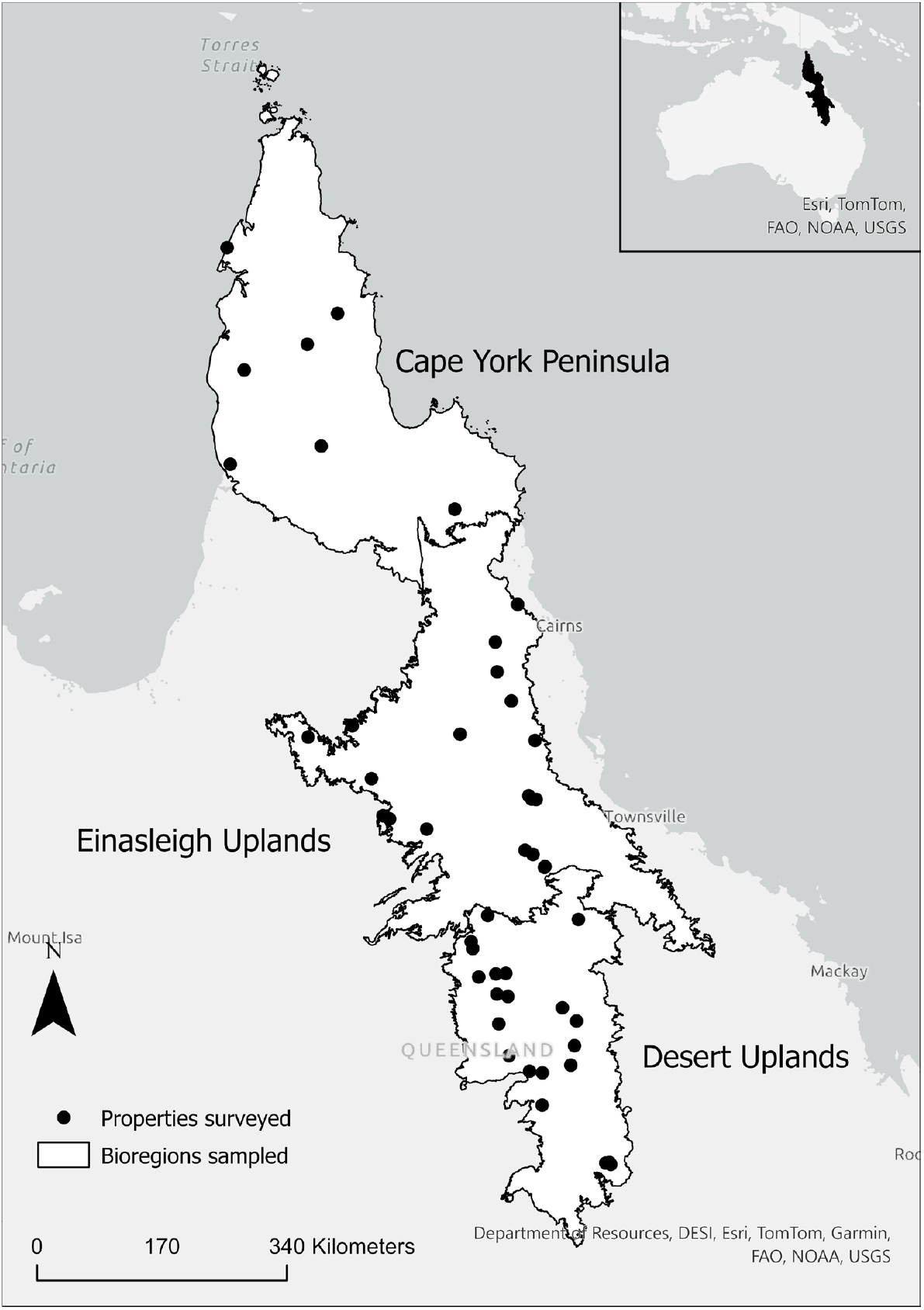
The location of the survey sites across the Desert Uplands, Einasleigh Uplands, and Cape York Peninsula. The black circles represent a group of sites on a single property.

The mammal data was collected from a standardised 1-ha fauna survey sites that sample for multiple terrestrial vertebrate groups (see details in Kutt and Fisher 2011). The mammal sampling comprised 20 Elliott traps (150 x 150 x 450 mm), two wire cage traps (300 mm x 300 mm x 600 mm), and four pitfall traps (600 mm deep x 300 mm wide) with drift fence. Elliott and cage traps were baited with either a peanut butter and oats ball, or meat flavoured dog biscuits. All traps were checked in the morning and afternoon and opened for five days / four nights. Trapping was supplemented by timed searches: three diurnal and two nocturnal searches each of 20 search-minutes duration conducted within the 1-ha site. The data was collected between 1998 and 2012.

A total of 240 sites were sampled in the Desert Uplands (mean site elevation 360.2 ± 6.5 m), 303 in the Einasleigh Uplands (mean site elevation 499.5 ± 11.8 m), and 182 in Cape York Peninsula (mean site elevation 97.1 ± 5.1 m). For each site, we calculated the total species richness and abundance for five mammal groups (excluding introduced species): all mammals combined, critical weight range mammals (mass <5.5kg) (Johnson and Isaac 2009), macropods, dasyurids and rodents, recognising there is overlap in species membership in some of the groups (Supplementary Material 1). Species richness is the total number of species (for each group) recorded by the standardised methods for a site, and abundance is the cumulative total of all captures and observations (for each group) recorded by the standardised methods for a site. We did not investigate single species models as there were very few species that had sufficient data to be modelled separately, and the distribution of species varied from bioregion to bioregion (i.e., naturally present in some, absent in others). In this research note, we were only concerned with broad patterns in species groups.

We derived three environmental variables for each survey site. These were (i) the average fractional cover green (i.e., green foliage and ground cover, hereafter referred to as Green) for the preceding 12-months to the survey, within a 500 m buffer (80 hectares) around the survey site (for further details on the derivation of fractional cover see Kutt *et al*. 2025), (ii) the elevation of the survey site derived from survey location using the 3 second (∼90m) Smoothed Digital Elevation Model Version 1.0 for Queensland (The State of Queensland 2025) (hereafter Elevation), and (iii) the vegetation diversity which was the number of mapped regional ecosystems (Neldner *et al*. 2012) within a 500 m buffer around the survey site (hereafter Veg Diversity). We chose these variables for several reasons, (a) fractional cover green reflects antecedent rainfall (Guerschman *et al*. 2020), which can determine mammal abundance (Lupone *et al*.), (b) elevation is not considered a feature of tropical savannas in Australia, yet in Queensland, there is a significant degree of elevation in the tropical savannas (Vanderduys *et al*. 2012) that could influence mammal pattern, and (c) tropical savanna woodlands are structurally similar across the three bioregions, though they change in dominant canopy vegetation from north to south. The use of vegetation type (i.e., regional ecosystem) as a predictor would simply reflect change in floristic dominance; therefore, we use vegetation diversity as this reflects heterogeneity or homogeneity of a site, irrespective of the vegetation type, aspects important for mammals in tropical savannas (Price *et al*. 2010).

### Analysis

We investigated the relationship between mammal abundance and species richness for each group and the three environmental variables collectively across the 725 sites using an information–theoretic model selection and averaging framework. The environmental data was first standardised to a 0–1 range using a min–max transformation to allow direct comparison of the strength of the estimates (Kutt *et al*. 2016). For each mammal group, we fitted generalized linear mixed-effects models with a negative binomial error distribution to account for overdispersion and the high frequency of zero counts. A global model including all three predictors was specified, and all possible subsets were generated using model dredging. Models were ranked according to the Akaike Information Criterion corrected for small sample size (AICc). Models with ΔAICc < 2 were retained as the best-supported candidate set, following the convention that such models are statistically indistinguishable in their level of empirical support (Burnham and Anderson 2002).

Where multiple candidate models were supported, we performed model averaging to estimate regression coefficients, unconditional standard errors, 95% confidence intervals for each predictor. Akaike weights were used to calculate the relative support for each model, and the sum of weights across all models containing a predictor was taken as a measure of its relative importance. A high cumulative weight (e.g. ≥0.8) was interpreted as strong support that a predictor was influential in explaining variation in mammal abundance, whereas lower weights indicated weaker evidence for predictor effects. All analyses were conducted in R version 4.4.3 (R Core Team 2024). Negative binomial models were fitted using the *MASS* package (Venables and Ripley 2002) and model selection and averaging were carried out with the *MuMIn* package (Bartoń 2024). Full model details are provided in Supplementary Material 2.

## RESULTS

A total of 46 native mammals from nine families were recorded across all three bioregions (Supplementary Material 1). In the Desert Uplands 23 species were recorded, in the Einasleigh Uplands, 33 species, and in Cape York Peninsula, only 18 species. The mammal species, and their mean abundance and site frequency in each bioregion is presented in Supplementary Material 1.

The mammal group models ranged from one to three terms (in addition to the intercept and negative binomial dispersion parameter) for the top two models (Table 1). For the best supported models, five had a single term (elevation and veg diversity twice, green once), four had two terms (green and veg diversity three times, elevation and green once) and one had all three terms (dasyurid abundance) (Table 1).

**Table 1.** The two best supported AIC models for the mammal groups. K is the number of model terms (including the intercept and the negative-binomial dispersion parameter). LogLik is the log likelihood, AICc is the Akaike Information Criterion corrected for small sample size, and Delta AIC is the relative difference between the AIC of one model and the best (lowest) AIC in all the models tested.

| Groups | Model | K | LogLik | AICc | DeltaAIC |
| --- | --- | --- | --- | --- | --- |
| Mammal abundance | ~ Green + Veg diversity | 4 | -1758.44 | 3524.93 | 0 |
| Mammal abundance | ~ Elevation + Green + Veg diversity | 5 | -1758.11 | 3526.30 | 1.37 |
| Mammal richness | ~ Elevation + Green | 4 | -1097.42 | 2202.90 | 0 |
| Mammal richness | ~ Elevation + Green + Veg diversity | 5 | -1096.53 | 2203.15 | 0.25 |
| Critical Weight Range abundance | ~ Veg diversity | 3 | -1433.67 | 2873.38 | 0 |
| Critical Weight Range abundance | ~ 1 | 2 | -1435.39 | 2874.79 | 1.41 |
| Critical Weight Range richness | ~ Elevation | 3 | -1026.13 | 2058.29 | 0 |
| Critical Weight Range richness | ~ Elevation + Veg diversity | 4 | -1025.36 | 2058.78 | 0.48 |
| Macropod abundance | ~ Green + Veg diversity | 4 | -1150.21 | 2308.47 | 0 |
| Macropod abundance | ~ Green | 3 | -1151.25 | 2308.54 | 0.07 |
| Macropod richness | ~ Green | 3 | -669.81 | 1345.66 | 0 |
| Macropod richness | ~ Elevation + Green | 4 | -669.32 | 1346.69 | 1.03 |
| Dasyurid abundance | ~ Elevation + Green + Veg diversity | 5 | -469.66 | 949.40 | 0 |
| Dasyurid abundance | ~ Elevation + Veg diversity | 4 | -472.11 | 952.27 | 2.87 |
| Dasyurid richness | ~ Green + Veg diversity | 4 | -332.72 | 673.50 | 0 |
| Dasyurid richness | ~ Elevation + Green + Veg diversity | 5 | -332.34 | 674.77 | 1.26 |
| Rodent abundance | ~ Elevation | 3 | -937.91 | 1881.84 | 0 |
| Rodent abundance | ~ Elevation + Green + Veg diversity | 5 | -936.03 | 1882.14 | 0.3 |
| Rodent richness | ~ Veg diversity | 3 | -579.57 | 1165.18 | 0 |
| Rodent richness | ~ Green + Veg diversity | 4 | -578.89 | 1165.83 | 0.65 |

The model averaging provided more insight to the key environmental parameters for each mammal group, including the size and direction of the effect (Figure 2). Vegetation diversity was positively associated with increasing mammal abundance, rodent and dasyurid species richness. Elevation was positively associated with increasing mammal and critical weight range species richness and decreasing rodent species richness and dasyurid abundance (Fig. 2). Fractional green cover was negatively associated with total mammal, macropod, and dasyurid abundance and richness (Fig. 2).

**Figure 2.**
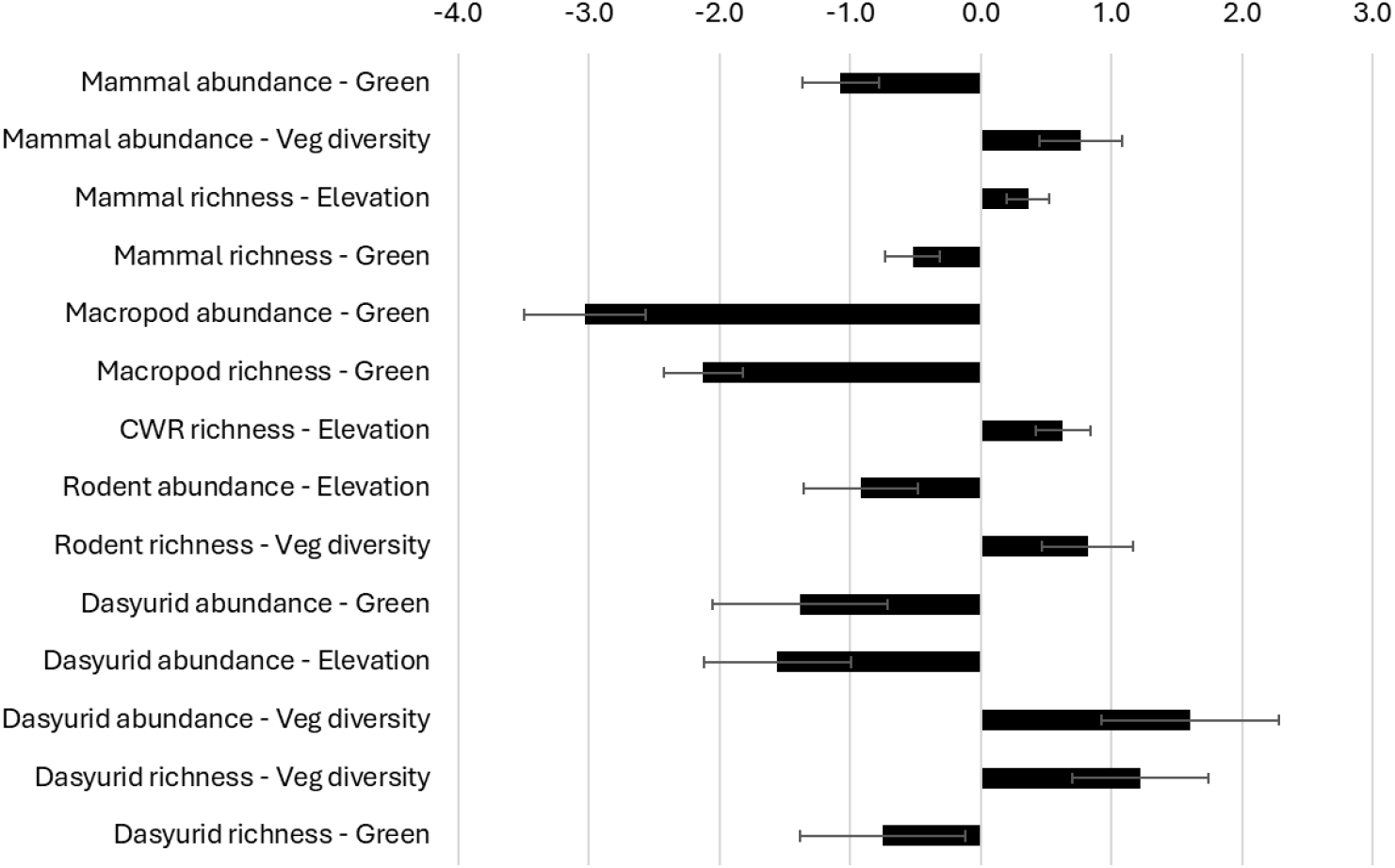
The significant parameter estimates and standard error the mammal groups identified via model averaging.

## DISCUSSION

In this study we examined a large systematic mammal survey data set (725 sites, sampled from 1998-2012) across a vast tropical savanna gradient to see if we could glean any coarse landscape scale patterns that might have some implications for conservation of mammal groups in northern Australia. We found that there were some clear relationships between the mammal groups and our three environmental parameters, and that there was variation between some groups (i.e., rodents and dasyurids) with some seemingly counterintuitive associations (i.e., negative effects with fractional cover green which represents tree cover and ground cover). This reinforces global analysis of mammal conservation that suggest that strategies and priorities need to consider taxonomic and regional factors to maximise protection (e.g., Brum *et al*. 2017).

We identify several limitations in our data and results. Firstly, we recognise that our data, despite the large number of sites, were comprised of many sites with few or no mammal data. We tried to account for this by looking at groups rather than species. Secondly, we use only three high level environmental predictor variables, rather than site-based habitat data. This may have missed nuanced patterns; however, we wanted to examine broad landscape scale patterns and elevation, green cover and vegetation diversity are all key drivers of wildlife pattern in northern Australia (Price *et al*. 2005). Lastly, our data did not include camera trapping, as some of the survey data pre-dated the wide use of camera trapping; this may reduce how comprehensive the site level mammal data is, and therefore interpretation.

The relationship of some mammal groups to elevation seems to have the most significant conservation implications, given in the past, tropical savannas have been considered topographically featureless (Woinarski *et al*. 2005). In Queensland, tropical savannas ranges from coastal plains just above sea level to higher altitude ecosystems above 1000 m (Vanderduys *et al*. 2012). Across the three bioregions we examined, the mean elevation for the sites sampled were 360 m (Desert Uplands), 500 m (Einasleigh Uplands) and 100 m (Cape York Peninsula). Both total mammal richness and critical weigh range mammal species richness was associated with increasing elevation; sites throughout the Einasleigh Uplands, that sit on the Great Dividing Range contain a diverse and sometime very abundant range of mammals including abundant populations of Greater glider *Petauroides volans*, Rufous bettong *Aepyprymnus rufescens* and Spectacled hare-wallaby *Lagorchestes conspicillatus* (DERM 2009). Conversely dasyurids and rodents identified a negative relationship with elevation; rodent species such as Canefield rat *Rattus sordidus* and Pale field-rat *Rattus tunneyi* occur in high abundance on the coastal floodplains of Cape York Peninsula (Perry *et al*. 2015), and genus such as *Planigale* and *Sminthopsis* are more commonly recorded in the semi-arid tropical savannas in the Desert Uplands, of intermediate to low altitude (Kutt 2004).

Vegetation cover (in our instance fraction cover green representing canopy and ground cover) is often associated with higher mammal abundance (Bergstrom *et al*. 2023). However megaherbivores in tropical savannas can suppress vegetation and wildlife (Wells *et al*. 2025), which in Australian tropical savannas comprise of domestic stock. The interaction between livestock, native herbivores (i.e., macropods) and rainfall, can cause complex interactions, not the least an increase in grazing pressure and a decline in ground cover (Witte and Croft 2025). Our analysis indicated macropods and dasyurids were negatively associated with green cover; however, both these groups are more species rich in the southern savannas where vegetation cover is lower, both naturally and compounded by the grazing of domestic stock (Kutt and Fisher 2011). In contrast, vegetation diversity, which is a coarse measure of landscape heterogeneity, had a positive relationship with mammal abundance overall, and rodents and dasyurids; previous work in Queensland found that tropical savannas containing a high diversity of grass, tree and bare ground cover contain more mammals (Price *et al*. 2010), and this seems to be the case in our results as well.

In this short study we examined a large set of systematically surveyed sites in the tropical savannas of Queensland to investigate broad landscape patterns in the mammal fauna. Though a reasonable number of species were recorded overall, the capture rate across sites was very low, but in keeping with many areas of the Queensland and the Top End tropical savannas, where there has been, respectively, an assumed decline in abundance (Kutt *et al*. 2023) and a well-documented and disquieting contemporary depletion (Woinarski *et al*. 2010). Our models provided some simple but important conclusions for the future conservation of mammals in north-eastern Queensland, namely (i) the higher altitude areas, which lie on the Great Dividing Range, are critical locations for smaller mammals, and deserve more focus for management, and future protection, especially in light of a changing climate, (ii) areas of higher vegetation diversity are more important for some taxa, and protection should not focus on elements perceived to offer high value (i.e., riparian areas) without the inclusion of surrounding areas of different vegetation types (Araújo *et al*. 2023). This has implications for fire and grazing management which protect riparian areas above the more extensive adjacent savannas (Lankester *et al*. 2009), and (iii) management that maintains vegetation heterogeneity (which respect to vegetation, ground cover and fire) will be important (Erdős *et al*. 2018).

## ACKNOWLEDGEMENTS

The data used in this study have been derived from a wide range of field surveys funded by a variety of Queensland and Australian Government programs up until 2012. Surveys were conducted under the conditions of Nature Conservation Act Scientific Purposes Permit NO/001480/96/SAA and NO/001480/99/SAA (EPA) and 1037/1038, 1170, 1359, 1513 and 1682 (Queensland Government). The data were collected in concert with a range of colleagues and volunteers over the years from CSIRO and the Queensland Department of the Environment, Tourism, Science and Innovation.

---

**Supplementary Material 1.**
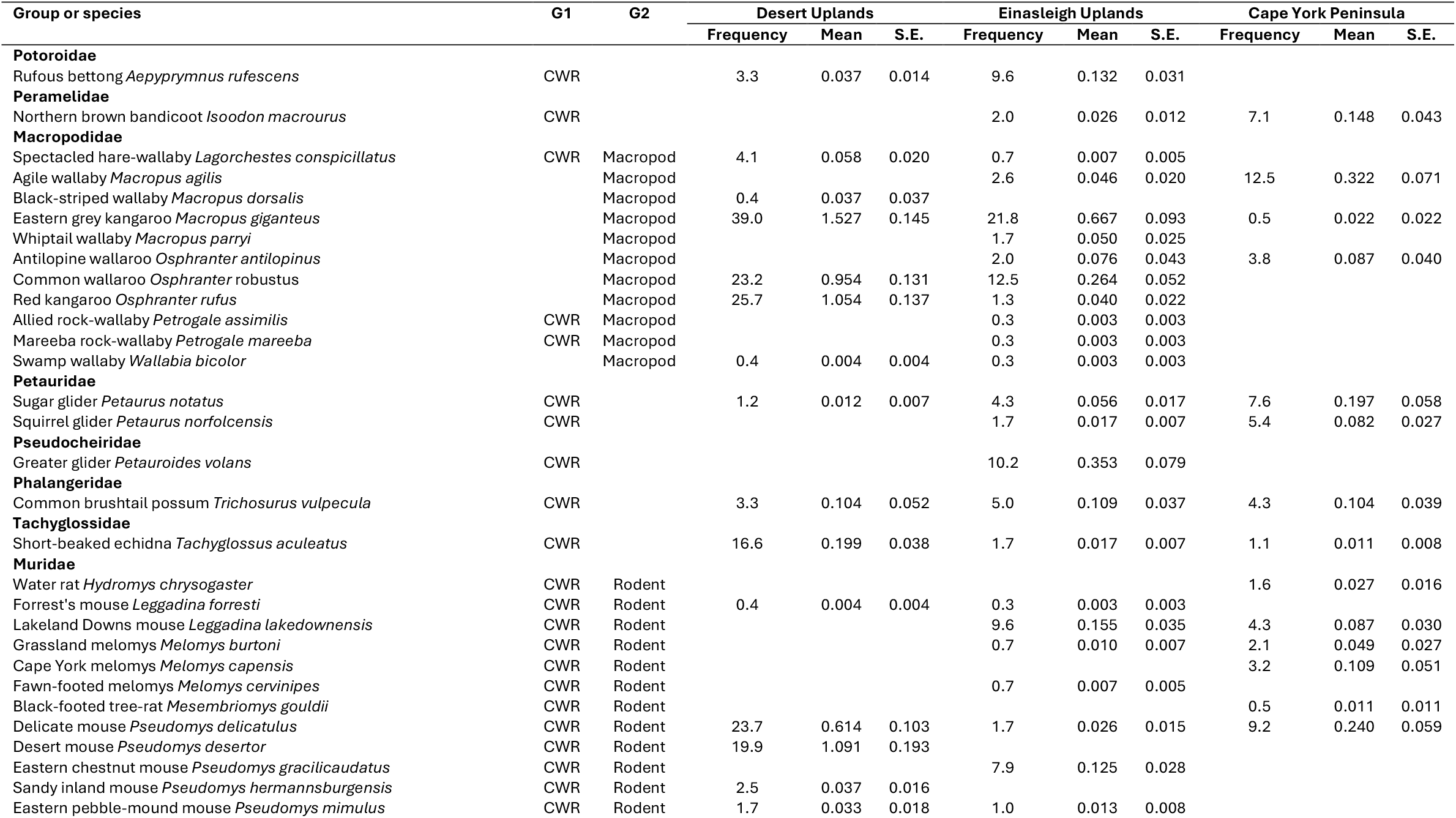
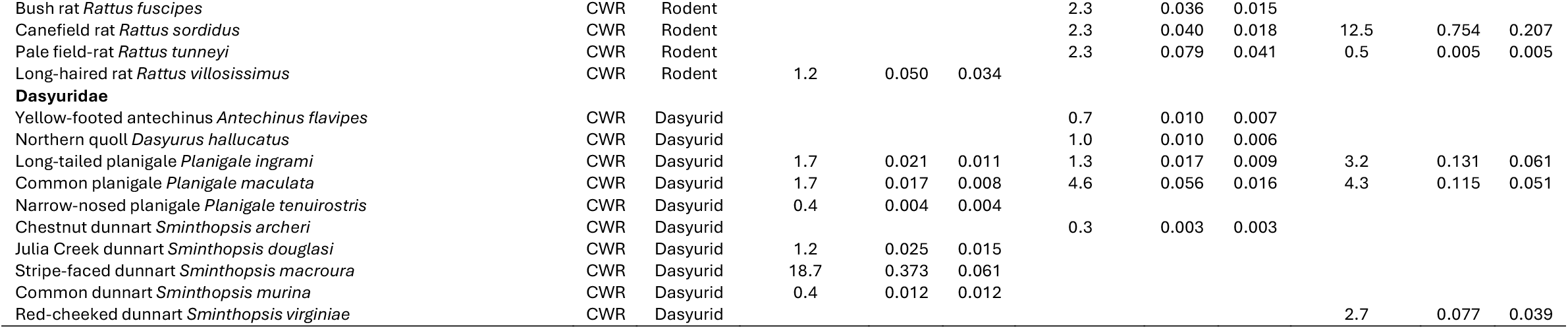
The mean (and standard error) of mammal species abundance recorded in each bioregion. Frequency is the percentage of sites a species was recorded, in that bioregion. G1 and G2 indicates mammal functional group. CWR = critical weight range.

**Supplementary Material 2.**
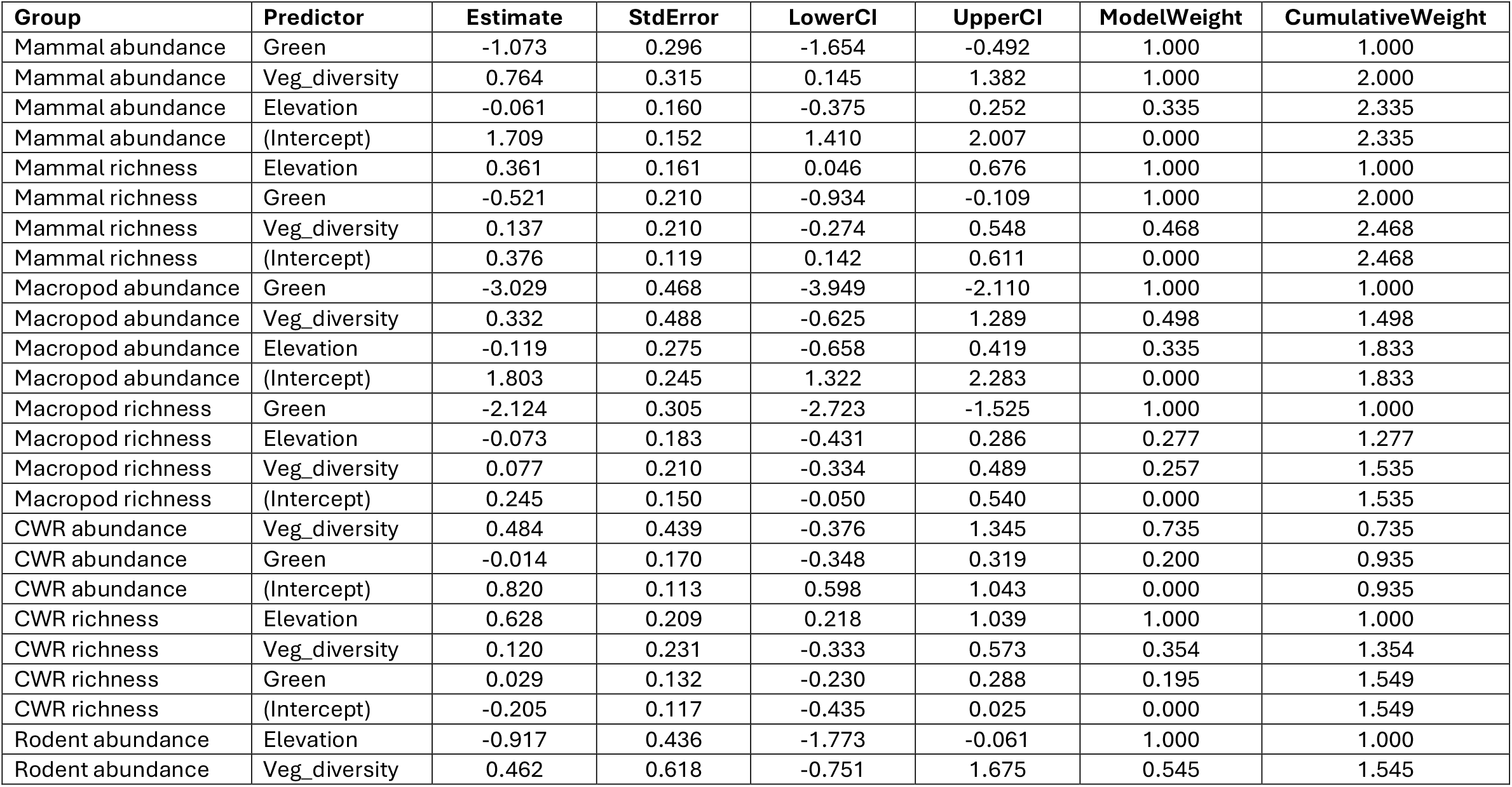
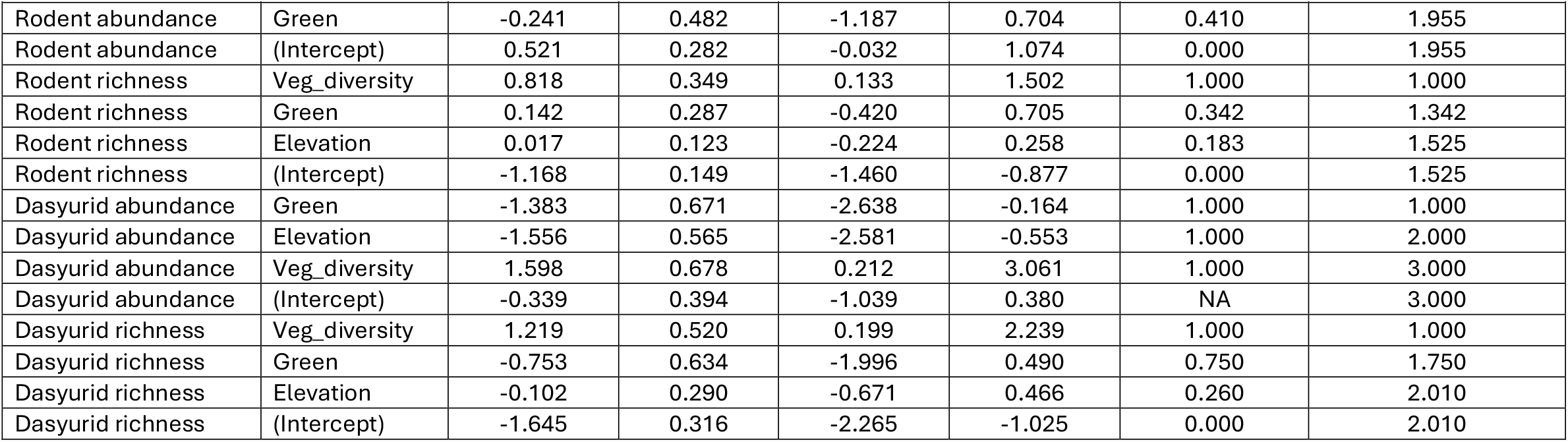
The results of the model averaging indicating the group, predictor, the estimate regression coefficients, unconditional standard errors, and 95% confidence intervals for each predictor. Akaike weights were used to calculate the relative support for each model, and the sum of weights across all models containing a predictor was taken as a measure of its relative importance. A high cumulative weight (e.g. ≥0.8) was interpreted as strong support that a predictor was influential in explaining variation in mammal abundance, whereas lower weights indicated weaker evidence for predictor effects.

| Group | Predictor | Estimate | StdError | LowerCI | UpperCI | ModelWeight | CumulativeWeight |
| --- | --- | --- | --- | --- | --- | --- | --- |
| Mammal abundance | Green | -1.073 | 0.296 | -1.654 | -0.492 | 1.000 | 1.000 |
| Mammal abundance | Veg_diversity | 0.764 | 0.315 | 0.145 | 1.382 | 1.000 | 2.000 |
| Mammal abundance | Elevation | -0.061 | 0.160 | -0.375 | 0.252 | 0.335 | 2.335 |
| Mammal abundance | (Intercept) | 1.709 | 0.152 | 1.410 | 2.007 | 0.000 | 2.335 |
| Mammal richness | Elevation | 0.361 | 0.161 | 0.046 | 0.676 | 1.000 | 1.000 |
| Mammal richness | Green | -0.521 | 0.210 | -0.934 | -0.109 | 1.000 | 2.000 |
| Mammal richness | Veg_diversity | 0.137 | 0.210 | -0.274 | 0.548 | 0.468 | 2.468 |
| Mammal richness | (Intercept) | 0.376 | 0.119 | 0.142 | 0.611 | 0.000 | 2.468 |
| Macropod abundance | Green | -3.029 | 0.468 | -3.949 | -2.110 | 1.000 | 1.000 |
| Macropod abundance | Veg_diversity | 0.332 | 0.488 | -0.625 | 1.289 | 0.498 | 1.498 |
| Macropod abundance | Elevation | -0.119 | 0.275 | -0.658 | 0.419 | 0.335 | 1.833 |
| Macropod abundance | (Intercept) | 1.803 | 0.245 | 1.322 | 2.283 | 0.000 | 1.833 |
| Macropod richness | Green | -2.124 | 0.305 | -2.723 | -1.525 | 1.000 | 1.000 |
| Macropod richness | Elevation | -0.073 | 0.183 | -0.431 | 0.286 | 0.277 | 1.277 |
| Macropod richness | Veg_diversity | 0.077 | 0.210 | -0.334 | 0.489 | 0.257 | 1.535 |
| Macropod richness | (Intercept) | 0.245 | 0.150 | -0.050 | 0.540 | 0.000 | 1.535 |
| CWR abundance | Veg_diversity | 0.484 | 0.439 | -0.376 | 1.345 | 0.735 | 0.735 |
| CWR abundance | Green | -0.014 | 0.170 | -0.348 | 0.319 | 0.200 | 0.935 |
| CWR abundance | (Intercept) | 0.820 | 0.113 | 0.598 | 1.043 | 0.000 | 0.935 |
| CWR richness | Elevation | 0.628 | 0.209 | 0.218 | 1.039 | 1.000 | 1.000 |
| CWR richness | Veg_diversity | 0.120 | 0.231 | -0.333 | 0.573 | 0.354 | 1.354 |
| CWR richness | Green | 0.029 | 0.132 | -0.230 | 0.288 | 0.195 | 1.549 |
| CWR richness | (Intercept) | -0.205 | 0.117 | -0.435 | 0.025 | 0.000 | 1.549 |
| Rodent abundance | Elevation | -0.917 | 0.436 | -1.773 | -0.061 | 1.000 | 1.000 |
| Rodent abundance | Veg_diversity | 0.462 | 0.618 | -0.751 | 1.675 | 0.545 | 1.545 |
| Rodent abundance | Green | -0.241 | 0.482 | -1.187 | 0.704 | 0.410 | 1.955 |
| Rodent abundance | (Intercept) | 0.521 | 0.282 | -0.032 | 1.074 | 0.000 | 1.955 |
| Rodent richness | Veg_diversity | 0.818 | 0.349 | 0.133 | 1.502 | 1.000 | 1.000 |
| Rodent richness | Green | 0.142 | 0.287 | -0.420 | 0.705 | 0.342 | 1.342 |
| Rodent richness | Elevation | 0.017 | 0.123 | -0.224 | 0.258 | 0.183 | 1.525 |
| Rodent richness | (Intercept) | -1.168 | 0.149 | -1.460 | -0.877 | 0.000 | 1.525 |
| Dasyurid abundance | Green | -1.383 | 0.671 | -2.638 | -0.164 | 1.000 | 1.000 |
| Dasyurid abundance | Elevation | -1.556 | 0.565 | -2.581 | -0.553 | 1.000 | 2.000 |
| Dasyurid abundance | Veg_diversity | 1.598 | 0.678 | 0.212 | 3.061 | 1.000 | 3.000 |
| Dasyurid abundance | (Intercept) | -0.339 | 0.394 | -1.039 | 0.380 | NA | 3.000 |
| Dasyurid richness | Veg_diversity | 1.219 | 0.520 | 0.199 | 2.239 | 1.000 | 1.000 |
| Dasyurid richness | Green | -0.753 | 0.634 | -1.996 | 0.490 | 0.750 | 1.750 |
| Dasyurid richness | Elevation | -0.102 | 0.290 | -0.671 | 0.466 | 0.260 | 2.010 |
| Dasyurid richness | (Intercept) | -1.645 | 0.316 | -2.265 | -1.025 | 0.000 | 2.010 |

## REFERENCES

Araújo R. P. d. C., Galliez M. & Bergallo H. G. (2023) Are riparian habitats always more diverse than nonriparian? A case study with small mammals in a rainforest environment. Canadian Journal of Zoology 101, 569–78.

Bartoń K. (2024) MuMIn: Multi-Model Inference_. R package version 1.48.4, <https://CRAN.R-project.org/package=MuMIn>.

Bergstrom B. J., Scruggs S. B. & Vieira E. M. (2023) Tropical savanna small mammals respond to loss of cover following disturbance: A global review of field studies. Frontiers in Ecology and Evolution Volume 11 -2023.

Brewster R., Jameson T., Roncolato F., Crowther M. S., Finnerty P. B. & Newsome T. M. (2024) Islands in the sky – could complex topography help us rewild beyond the fence? Pacific Conservation Biology 30, -.

Brum F. T., Graham C. H., Costa G. C., Hedges S. B., Penone C., Radeloff V. C., Rondinini C., Loyola R. & Davidson A. D. (2017) Global priorities for conservation across multiple dimensions of mammalian diversity. Proceedings of the National Academy of Sciences 114, 7641–6.

Burnham K. P. & Anderson D. R. (2002) Model selection and multimodel inference: a practical information-theoretic approach. Springer-Verlag, New York, USA.

DERM. (2009) Biodiversity Planning Assessment, Einasleigh Uplands Bioregion Fauna Expert Panel Report, Version 1.1. Department of Environment and Resource Management, Queensland Government, Brisbane.

Doherty T. S., Watchorn D. J., Miritis V., Pestell A. J. & Geary W. L. (2023) Cats, foxes and fire: quantitative review reveals that invasive predator activity is most likely to increase shortly after fire. Fire Ecology 19, 22.

Erdős L., Kröel-Dulay G., Bátori Z., Kovács B., Németh C., Kiss P. J. & Tölgyesi C. (2018) Habitat heterogeneity as a key to high conservation value in forest-grassland mosaics. Biological Conservation 226, 72–80.

Guerschman J. P., Hill M. J., Leys J. & Heidenreich S. (2020) Vegetation cover dependence on accumulated antecedent precipitation in Australia: Relationships with photosynthetic and non-photosynthetic vegetation fractions. Remote Sensing of Environment. 240, 111670.

Johnson C. N. & Isaac J. L. (2009) Body mass and extinction risk in Australian marsupials: The ‘Critical Weight Range’ revisited. Austral Ecology 34, 35–40.

Kutt A. S. (2004) Patterns in the composition and distribution of the vertebrate fauna, Desert Uplands Bioregion, Queensland. School of Marine and Tropical Biology, James Cook University of North Queensland, Townsville.

Kutt A. S. & Fisher A. (2011) Increased grazing and dominance of an exotic pasture (Bothriochloa pertusa) affects vertebrate fauna species composition, abundance and habitat in savanna woodland. The Rangeland Journal 33, 49–58.

Kutt A. S., Healy A. J. & Hamer R. P. (2025) Birdseye in the sky: the relationship between fractional cover, rainfall and woodland birds in a partially modified tropical savanna. The Rangelands Journal 4.

Kutt A. S., Vanderduys E. P., Perry J. J., Mathieson M. T. & Eyre T. J. (2016) Yellow-throated miners Manorina flavigula homogenize bird communities across intact and fragmented landscapes. Austral Ecology 41, 316–27.

Kutt A. S., Waller N. L., Colman N. J., Perry J. J. & Starr C. R. (2023) Camera trapping ekes out some improvement for surveying sparse mammal populations in northern Queensland. Australian Mammalogy 45, 293–304.

Lankester A., Valentine P. & Cottrell A. (2009) ‘The sweeter country’: social dimensions to riparian management in the Burdekin rangelands, Queensland. Australasian Journal of Environmental Management 16, 94–102.

Legge S., Woinarski J. C. Z., Burbidge A. A., Palmer R., Ringma J., Radford J. Q., Mitchell N., Bode M., Wintle B., Baseler M., Bentley J., Copley P., Dexter N., Dickman C. R., Gillespie G. R., Hill B., Johnson C. N., Latch P., Letnic M., Manning A., McCreless E. E., Menkhorst P., Morris K., Moseby K., Page M., Pannell D. & Tuft K. (2018) Havens for threatened Australian mammals: the contributions of fenced areas and offshore islands to the protection of mammal species susceptible to introduced predators. Wildlife Research 45, 627–44.

Lupone L., Cooke R., Rendall A. R., Siegrist A., Penton C., Carlyon M., Ouchtomsky T. & White J. G. Hindcasting long-term data unveils the influence of a changing climate on small mammal communities. Divers. Distrib. n/a, e13901.

Neldner V. J., Wilson B. A., Thompson E. J. & Dillewaard H. A. (2012) Methodology for Survey and Mapping of Regional Ecosystems and Vegetation Communities in Queensland. Version 3.2. Updated August 2012. p. 124, Brisbane.

Perry J. J., Vanderduys E. P. & Kutt A. S. (2015) More famine than feast: pattern and variation in a potentially degenerating mammal fauna on Cape York Peninsula. Wildlife Research 42, 475–87.

Price B., Kutt A. S. & McAlpine C. A. (2010) The importance of fine-scale savanna heterogeneity for reptiles and small mammals. Biological Conservation 143, 2504–13.

Price O., Rankmore B., Milne D., Brock C., Tynan C., Kean L. & Roeger L. (2005) Regional patterns of mammal abundance and their relationship to landscape variables in eucalypt woodlands near Darwin, northern Australia. Wildlife Research 32, 435–46.

R Core Team. (2024) R: A language and environment for statistical computing. R Foundation for Statistical Computing, Vienna, Austria. https://www.R-project.org/.

The State of Queensland. (2025) Digital Elevation Model -3 second -Queensland. The Department of Natural Resources and Mines, Manufacturing and Regional and Rural Development, Brisbane.

Vanderduys E. P., Kutt A. S. & Kemp J. E. (2012) Upland savannas: the vertebrate fauna of largely unknown but significant habitat in north-eastern Queensland. Australian Zoologist 36, 59–74.

Venables W. N. & Ripley B. D. (2002) Modern applied statistics with S. Fourth Edition, Springer, New York; Berlin.

Wallach A. D. & Lundgren E. J. (2025) Review of evidence that foxes and cats cause extinctions of Australia’s endemic mammals. BioScience.

Wells H. B. M., Kimuyu D. M., Veblen K. E. & Young T. P. (2025) Megaherbivores suppress precipitation-driven plant irruptions in a tropical savanna. Ecosphere 16, e70239.

Witte I. & Croft D. B. (2025) Who Eats the Grass? Grazing Pressure and Interactions Between Wild Kangaroos, Feral Goats and Rabbits, and Domestic Sheep on an Arid Australian Rangeland. Wild 2, 5.

Woinarski J. C. Z., Armstrong M., Brennan K., Fisher A., Griffiths A. D., Hill B., Milne D. J., Palmer C., Ward S., Watson M., Winderlich S. & Young S. (2010) Monitoring indicates rapid and severe decline of native small mammals in Kakadu National Park, northern Australia. Wildlife Research 37, 116–26.

Woinarski J. C. Z., Fisher A. & Milne D. (1999) Distribution patterns of vertebrates in relation to an extensive rainfall gradient and variation in soil texture in the tropical savannas of the Northern Territory, Australia. Journal of Tropical Ecology 15, 381–98.

Woinarski J. C. Z., Williams R. J., Price O. & Rankmore B. (2005) Landscapes without boundaries: wildlife and their environments in northern Australia. Wildlife Research 32, 377–88.

